# Single-Cell Trajectory Inference for Detecting Transient Events in Biological Processes

**DOI:** 10.1101/2025.05.07.652753

**Authors:** Alexandre Hutton, Jesse G. Meyer

## Abstract

Transient surges in gene or protein expression often mark the key regulatory checkpoints that propel cells from one functional state to the next, yet they are easy to miss in sparse, noisy single-cell omics data. We introduce **scTransient**, a trajectory-inference pipeline integrated into our cloud-based single-cell analysis platform PSCS. scTransient transforms single-cell expression profiles into continuous pseudotime signals and couples them with wavelet-based signal processing to isolate short-lived but biologically meaningful bursts of activity. After ordering cells with unsupervised graph trajectories or supervised psupertime, **scTransient** windows expression values along pseudotime, applies a continuous wavelet transform, and assigns every gene a Transient-Event Score (TES) that rewards sharp, isolated coefficients while penalizing background fluctuations. Synthetic benchmarks show TES robustly recovers transient events across a wide range of cell numbers, signal-to-noise ratios, and event widths. Applying scTransient to three public datasets— hematopoietic differentiation, monocyte-to-macrophage maturation, and single-cell proteomic cell-cycle progression—uncovers previously unreported, process-specific expression spikes. These include erythropoiesis regulators (e.g., Nfe2), membrane-raft remodeling proteins during macrophage differentiation, and S-phase DNA-replication factors in A549 cells. Functional enrichment confirms that top-scoring genes cluster into pathways directly pertinent to each transition. By extending trajectory inference from descriptive ordering to quantitative detection of fleeting regulatory programs, scTransient—now readily accessible via the PSCS web interface—offers researchers a practical route to uncovering transient molecular events that drive development, differentiation, and disease.

## Introduction

Omic profiles can describe the functional state of individual cells. The quantification of mRNA expression or proteins is continuous by nature, supporting the idea that cell states are inherently not discrete and that they are better understood as a continuum [1], [2]. Cells nonetheless experience changes in their omic profiles as the cells transition between states, and some of these processes require transient events (TEs) that guide the cell’s molecular profile to match its intended function. We expect that for some processes, it is possible to detect these TEs by measuring the omic profiles of cells along that process. Specifically, we distinguish between long-term changes that reflect a change in a cell’s function and state-related changes that are only relevant as the cell transitions from one state to another. Since we do not know *a priori* which genes are relevant for which processes, methods permitting greater depth in omic profiles are appealing. Methods for acquiring such profiles are generally destructive of the sample, limiting our ability to track changes within a single cell. Trajectory inference is intended to relate single-cell samples to one another and establish a rough progression of state between samples in a dataset. Although trajectory inference methods are not inherently tied to time, it is possible to use it to examine transitions by connecting a trajectory to a known biological process (e.g., by looking at markers of cell cycle differentiation).

We introduce a method for detecting these TEs that are represented as temporary changes in gene expression by examining the pseudotimecourse of a biological process, summarized in **Fig. 1**. Identification is complicated by several factors: (1) single cell omics experiments can now quantify the products from thousands of genes, making false positives more likely, (2) ensuring that the pseudotimecourses relate to a particular process requires external process-related measurements, and (3) it is not known *a priori* what these transitions should look like in pseudotime.

**Figure 1.**
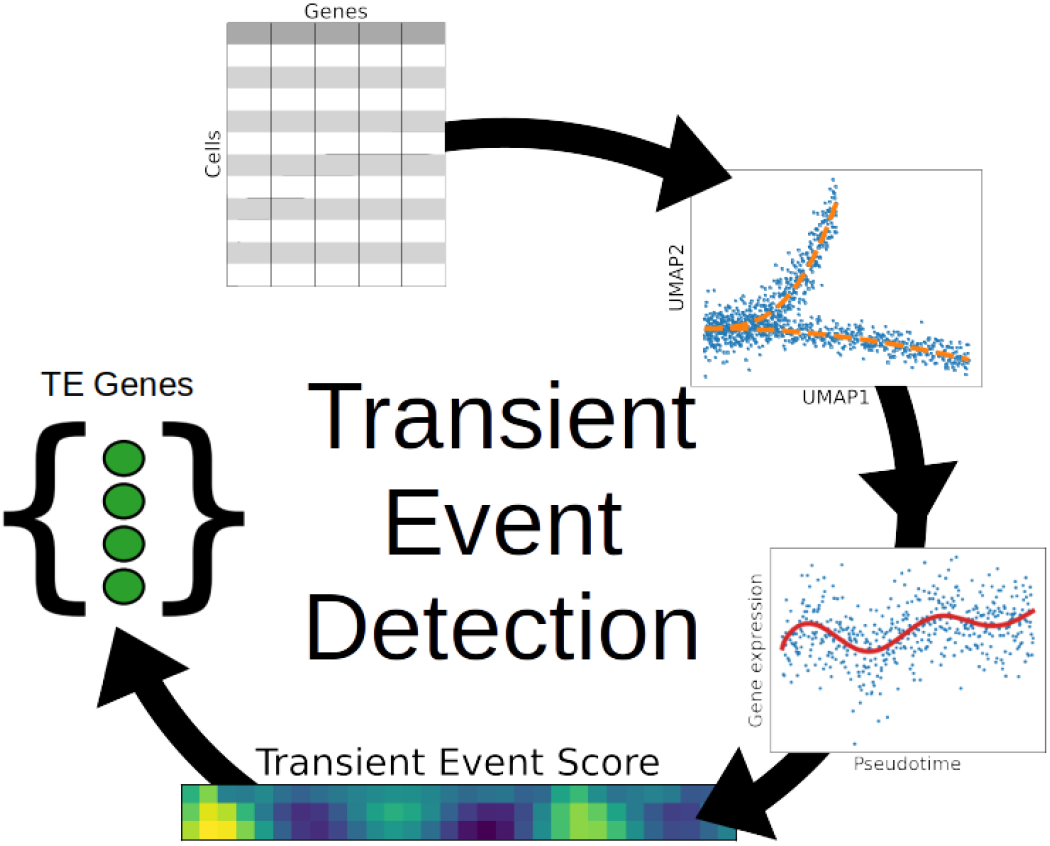
Schematic for detecting genes containing transient events. Starting with the gene quantifications, obtain a trajectory with pseudotime ordering. Convert samples along that trajectory to a pseudotime signal by windowing along pseudotime, then obtain the wavelet coefficients of the signal. From the coefficients, extract a set of genes by scoring the coefficients and selecting genes above a selected threshold.

We aim to solve these challenges as follows:

1. Compare the identified genes to known processes. We can validate the method by examining the set of genes we identify to those that are known to be involved in a particular process. However, we note that any method examining many timecourses is ultimately susceptible to false positives, particularly where the signal-to-noise ratio is low.
2. Use known cell biomarkers that establish the progression of the single-cell samples in pseudotime along a known biological process. For example, cell cycle markers to identify the phase of different samples or cell type markers to identify differentiation along pseudotime.
3. Use a wavelet transform to obtain a scale-adjustable function that can detect transient changes. By finding peaks in the coefficients of the wavelet transform, we can identify positions in pseudotime corresponding to the TEs.

## Methods

### Trajectory inference and pseudotimecourses

Trajectory inference is the process of linking single-cell samples to one another by examining their omic profiles and inferring an ordering to them. Since omic data acquisition of single-cell samples is destructive, it is necessary to combine the measurements of different cells together to imitate monitoring over the course of a biological process. For this paper, we use two approaches to map cell samples to a pseudotime axis. First, we combine PAGA [3] with Djikstra’s algorithm [4] and DPT [5] to obtain the subset of cells relevant to a specific biological process. Second, we use a supervised pseudotime method, *psupertime* [6], for processes with known and available biomarkers.

Once cells are ordered on a pseudotime axis, we can convert the collection of individual samples into a signal by using an averaging window along pseudotime. Windowing can be parameterized by the type, number, and width of the windows. Parameter selection at this stage can significantly impact the signal and consequently the information that can be reliably extracted from it: the type of window dictates the relative importance of different data points, the number of windows controls how finely the signal is represented, and the width of the window controls how many points are considered.

With the cells converted to a signal representing gene expression along a process, we can use signal processing to identify transient activity and localize it in pseudotime.

### Wavelet transform

Signals can be deconstructed and represented in different ways, with different utility. The most useful and common tool is the fast Fourier transform (FFT), which allows us to deconstruct discrete signals into sinusoidal signals and their phase. Modifications such as the short-time Fourier transform (STFT) allows for the FFT to be localized to part of the signal. Without knowing the width of the transient signal, we would need to apply the STFT at multiple window widths. This can be done more efficiently by using the continuous wavelet transform (CWT) [7], which incorporates the width and position of the subsignal, making it ideal for analyzing signals with uncertain properties. The CWT decomposes the signal into signals that are parameterized by their position (point in pseudotime) and scale (width of the signal). It produces coefficients for every combination of scale and position, and by identifying extrema in the coefficients we can use it to identify transient changes in gene expression.

### Transient event score

#### Gene expression signals

One major limitation for the described method is that the resulting gene expression signals can be noisy. Moreover, given that there can be thousands of quantified genes in an omic dataset, manually reviewing the wavelet transform would require too much manual intervention and lead to false positives. Instead, we use a heuristic to score all genes and retain only top-scoring genes. The heuristic, “transient event score” (TES), consists of two terms derived from the coefficients obtained from the wavelet transform of the pseudotime signal. We first compute the modified Z score of each coefficient *c* in the set of coefficients *C* (Eq. 1). We then multiply the coefficients with the modified Z score and take the maximum of the absolute value of the product (Eq. 2).

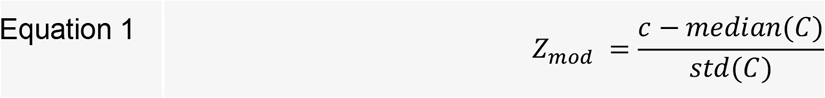

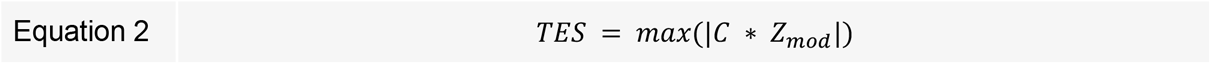

High-magnitude coefficients correspond to wavelets that match the signal better at that point, while the modified Z score selects for genes that have low variance outside of the spike (e.g., having multiple, smaller spikes would reduce the Z score for a gene, suggesting that the spikes could be noise).

### Synthetic data generation

Validating the proposed approach using existing datasets is difficult as there are multiple confounding factors. First, establishing a verifiable pseudotime ordering is difficult; linking an obtained pseudotime to a specific biological process when cells undergo multiple processes at a given time [8] requires sample metadata that is not always available. In cases where such metadata is available (e.g., cell type labeling), it is not always clear whether cell labels are sufficiently reliable; although it is useful to discretize cell state for certain applications including communication, the continuous nature of cellular omic profiles makes strict boundaries between states questionable. Second, we do not have ground truth for a given process and dataset. While a particular dataset might contain observations related to a particular process, it is possible that it does not capture the transient changes that the method is trying to identify. Failing to identify a change that is not present would then not be a failure of the method; similarly, identifying an expected change that is not actually present would not be a success. Third, knowing the likelihood of picking up false positives, it becomes easier to dismiss genes that appear odd and focus on identified genes that “make sense” for a particular dataset. This is particularly problematic as the intent of the method is to identify new genes that may not be known to fit at the time of analysis.

To address some of these confounds, we start by evaluating the method on synthetic data. For any dataset, we can dictate the pseudotime ordering to be correct and related to a process of interest. It also allows us to set which genes contain a valid transient signal, along with exploring the effect of different processing steps on downstream conclusions. Synthetic data also grants us the ability to evaluate the impact of different aspects of the dataset on performance metrics such as precision and recall. We emphasize that the utility of a synthetic dataset is limited by how well it represents real data. Even with a coarse correspondence between real and synthetic data, it is useful as a first-pass validation.

#### Synthetic data parameters

We parameterize the synthetic data as follows. We first set the total number of cells and genes. Pseudotime is assumed to be in the range of 0-1. Along pseudotime, we use a probability density function to control where in pseudotime samples are more likely to appear (e.g. a fast transition from one state to another could result in only a few samples in between two states). Genes are separated into two categories: either “signal” or “noise” genes. Signal genes contain the process-related transient change that the method is trying to detect; noise genes contain only noise. The signal genes contain a set number of spikes at locations in pseudotime, and each spike has a specified width and amplitude. Additive noise is included with all genes and consists of white noise with autocorrelation introduced by weighting the previous sample based on its distance in pseudotime.

## Preprocessing pipelines

The pipelines described here are available on [PSCS](https://pscs.xods.org/p/jqNzO) [9], a platform for producing reproducible analyses for single-cell experiments. Note that in our analyses, we kept zero values, but users of scTransient may prefer to exclude missing values.

### Hematopoiesis

The data consists of 2730 samples with 3451 genes. Functions made available through Scanpy [10] were used for preprocessing. We first computed the coverage of each gene, defined as the fraction of samples where the gene was detected. Genes with less than 50% coverage were dropped, resulting in 570 genes. Cell quantities were normalized, had log_e_(x+1) applied, and rescaled to 0 mean and 1 standard deviation. PCA was computed using the ARPACK solver [11]. Sample neighborhoods were calculated using n_neighbors=4, n_pcs=20.

Leiden clustering, the diffusion map, and DPT were then computed with default parameters. PAGA connectivities were then computed for the group labels supplied with the dataset (“paul15_clusters”).

### Monocyte to macrophage transition

The joint protein and transcriptomic data was downloaded from the SCoPE2 page [12] [link]. The data has already been scaled; PCA was computed using the arpack solver, neighborhoods were computed using n_neighbors=20 and n_pcs=40. Leiden clustering and DPT were computed with default parameters.

### Cell cycle

The dataset was produced by processing the raw data made available through the ProteomeXchange Consortium [13] via PRIDE [14] with ID PXD049412. Cells were filtered to have at least 2000 genes, and genes were required to be present in at least 3 cells. Normalization and log_e_ before regressing out the number of genes as a confound. Data was rescaled to 0 mean and 1 standard deviation with a value capped at 10.

PCA was computed using the arpack solver, sample neighborhoods were computed with n_neighbors=10 and n_pcs=40. Cell cycle phase labels were predicted by selecting known cell-cycle-related genes from [regev et al], assigning phase labels to each sample using Scanpy’s *score_genes_cell_cycle* function, then setting the phase order as G1=1, S=2, and G2M=3. Using these ordinal labels, *psupertime* was used to predict a pseudotime value for each value.

#### Code Availability

The scTransient pipeline is available on [PSCS](https://pscs.xods.org/p/jqNzO) [14] and on [GitHub](https://github.com/xomicsdatascience/scTransientNotebooks).

#### Data Availability

Datasets were downloaded from external sources: hematopoiesis [15] via Scanpy’s [10] datasets, SCoPE2 [12] via the project website, and the cell cycle dataset [16] is available through the ProteomeXchange Consortium [13] via PRIDE [14] with ID PXD049412.

## Results

We present two sets of results: results from synthetic data and results from existing public datasets characterizing different biological processes.

### Synthetic data results

Results from the synthetic datasets are intended to evaluate the method under known conditions. These serve as a sanity check to verify that the method functions as expected and to explore the impact of changes in the data on downstream analyses and conclusions.

The explicit assumption for this method is that there are detectable transient changes in gene expression. We explore some of the major effects present in single-cell experiments, including sample count, noisy samples, and sample density along pseudotime.

#### Sample count and SNR

We simulated datasets to investigate the effect of both sample count and the signal-to-noise ratio (SNR) on the detectability of transient events. The number of acquired cells has a number of impacts on the detectability of a signal. As demonstrated in **Figures 2A, 2B** the cellular resolution in pseudotime determines whether a transient event is detectable even when it is known to exist. More importantly, in the typical scenario with a thousand quantified genes, automatic detection above statistical thresholds is important. We generated datasets with multiple signal genes of varying SNR and sample counts (**Fig. 2C**). Examining the score, we see a steady increase in event score with increasing SNR. However, low-sample datasets (n={10,20}) also had an increasing variance in scoring. For the noise genes in those same datasets, we see a generally-decreasing trend in both the mean score and the score variance as the number of samples increases (**Fig. 2D**). For low-sample experiments, the score distribution is bimodal (**Fig. 2E**), where the higher scores are from instances where the pseudotime samples were located at the transient event. Although the likelihood of a false negative decreases with SNR, it reaches a minimum of 50-60% for 10 samples, 30-40% for 20 samples, and 0% for 50 samples with a sufficient SNR (**Fig. 2F**).

**Figure 2.**
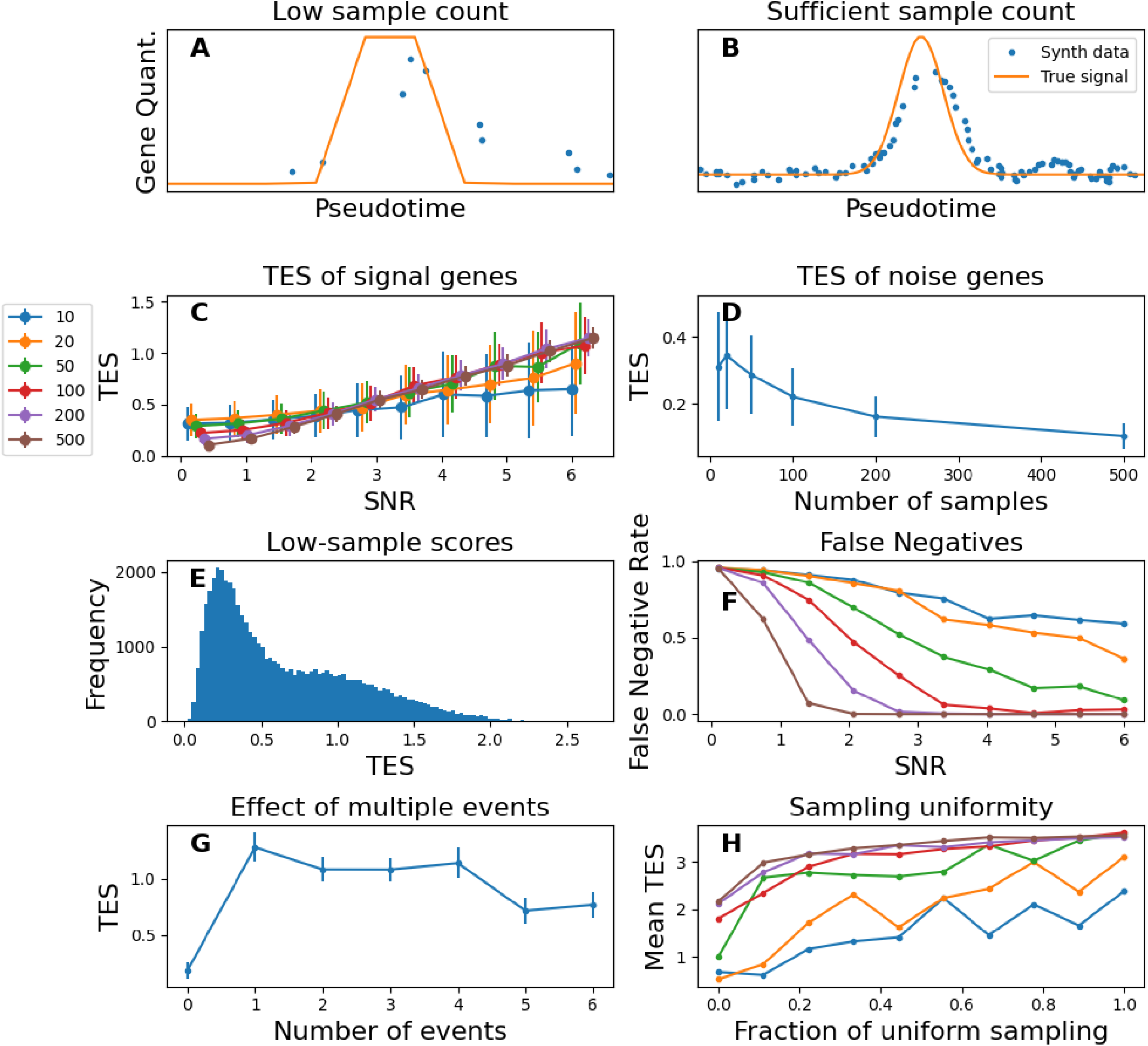
TRE detection with the synthetic datasets. **(A-B)** The number of cell samples in a dataset impacts whether a transient change in expression is observable. **(C)** Evaluation of the impact of the number of samples and the SNR on the score given to signal genes. **(D)** For a set SNR, the sample size directly affects the scores given to noise-only genes, establishing a noise floor for different conditions. **(E)** For multiple random instantiations of a dataset, the distribution of scores for low-sample datasets produces a bimodal distribution with higher scores being predicted when samples appear in the during transient expression change. **(F)** The rate of signal genes being labeled as noise genes. The score threshold is set as two standard deviations above the mean noise gene score for a dataset. **(G)** The effect of genes demonstrating multiple events in a single pseudotimecourse is to decrease the overall score. **(H)** Rapid cell state transitions are modeled as a decrease in the uniformity of sampling across pseudotime. Starting on the left with a reduction to no samples obtained at the peak of the transition and moving to the right to be completely uniform, low-sample counts are greatly impacted.

#### Multiple transient events

The proposed heuristic consists of two components: the coefficient magnitude and the modified Z score. We investigated the impact of the presence of multiple TEs on the overall score assigned to signal genes, reasoning that additional TEs would increase the standard deviation of coefficients and lower the overall score. The results are shown in **Fig. 2G**, where starting with the first event, we see a general decreasing trend in the gene score as the number of TEs increases.

#### Duration of transient event

The pseudotime axis represents a measure of change in the gene expression and does not have a direct correspondence to time. Despite that, the timing of sample acquisition has an important effect on data acquired. We model the speed in transitions by the density of cell samples; faster transitions have fewer samples in the area of pseudotime where the signal is present. The duration of the TE is particularly impactful in cases where there are only a few samples. We reduce the probability density proportional to the magnitude of the signal, making it less likely to acquire samples located at the peak of the TE signal. **Figure 2H** shows the impact of the sampling density for different sample counts, with 0 indicating that no samples are found at the signal’s peak, and 1 indicating that it is as likely as elsewhere in pseudotime (i.e., uniform distribution). We see a general increase for all sample counts as sampling becomes more uniform.

### Analysis of Existing Datasets

#### Hematopoiesis

Moving beyond synthetic datasets, we can analyze existing datasets to verify whether the proposed method identifies known biological processes. One dataset of interest is made widely available through Scanpy [10], from Paul et al. 2015 [15] quantifying the transcriptomic profiles of the multiple cell types present in hematopoiesis. Applying a preprocessing pipeline described in the supplemental materials, we obtain trajectories by computing PAGA connectivities (**Fig. 3A**), selecting relevant endpoints, and then using Djikstra’s algorithm to identify the progression of cell states. The trajectory can be seen in **Fig. 3B**, where it starts in the cluster identified as containing stem cells and proceeds through the megakaryocyte-erythroid progenitor cells, and then through different erythrocyte clusters. Using this trajectory, we order the cells based on DPT-derived pseudotimes, obtain pseudotimecourses, and identify genes containing TEs using scTransient. The proportion of cell labels along pseudotime (**Fig. 3C**) show the expected shift from progenitor cell states to differentiated erythrocytes. Scoring the available genes and showing their distribution in **Fig. 3D**, we apply a score cutoff of 4 and obtain 12 genes most likely to exhibit TEs: Alad, Blvrb, Car2, Fam132a, Fth1, Glrx5, Gnb2l1, Nedd4, Nfe2, Prdx2, Prdx6, Tmem14c. The pseudotimecourse for Nfe2 is shown in **Fig. 3E**, along with the scoring heuristic across wavelet scale and time. Nfe2 is a transcription factor required for erythrocyte function and maturation [17], [18], [19]. We performed gene enrichment analysis with g:Profiler [20], [21] using the set of genes and identified several GO terms relevant to erythrocytes (**Table 1**): peroxiredoxin activity, cell redox homeostasis, heme metabolic process.

**Table 1.** Functional gene enrichment pathways erythropoiesis pathway of the hematopoiesis dataset as identified by the proposed method. Bold terms indicate pathways that are particularly relevant to erythrocyte activity.

| source | native | name | p_value |
| --- | --- | --- | --- |
| GO:BP | GO:0045454 | **cell redox homeostasis** | 0.000687 |
| GO:MF | GO:0051920 | **peroxiredoxin activity** | 0.000799 |
| GO:BP | GO:0042168 | **heme metabolic process** | 0.000969 |
| GO:BP | GO:0006778 | **porphyrin-containing compound metabolic process** | 0.00148 |
| GO:BP | GO:0033013 | **tetrapyrrole metabolic process** | 0.002365 |
| GO:BP | GO:0042440 | pigment metabolic process | 0.005062 |
| GO:CC | GO:0070062 | extracellular exosome | 0.005995 |
| GO:CC | GO:1903561 | extracellular vesicle | 0.006397 |
| GO:CC | GO:0043230 | extracellular organelle | 0.006414 |
| GO:CC | GO:0065010 | extracellular membrane-bounded organelle | 0.006414 |
| GO:MF | GO:0016491 | **oxidoreductase activity** | 0.031954 |
| GO:MF | GO:0004601 | **peroxidase activity** | 0.043444 |
| GO:MF | GO:0016684 | **oxidoreductase activity, acting on peroxide as...** | 0.046611 |

**Figure 3.**
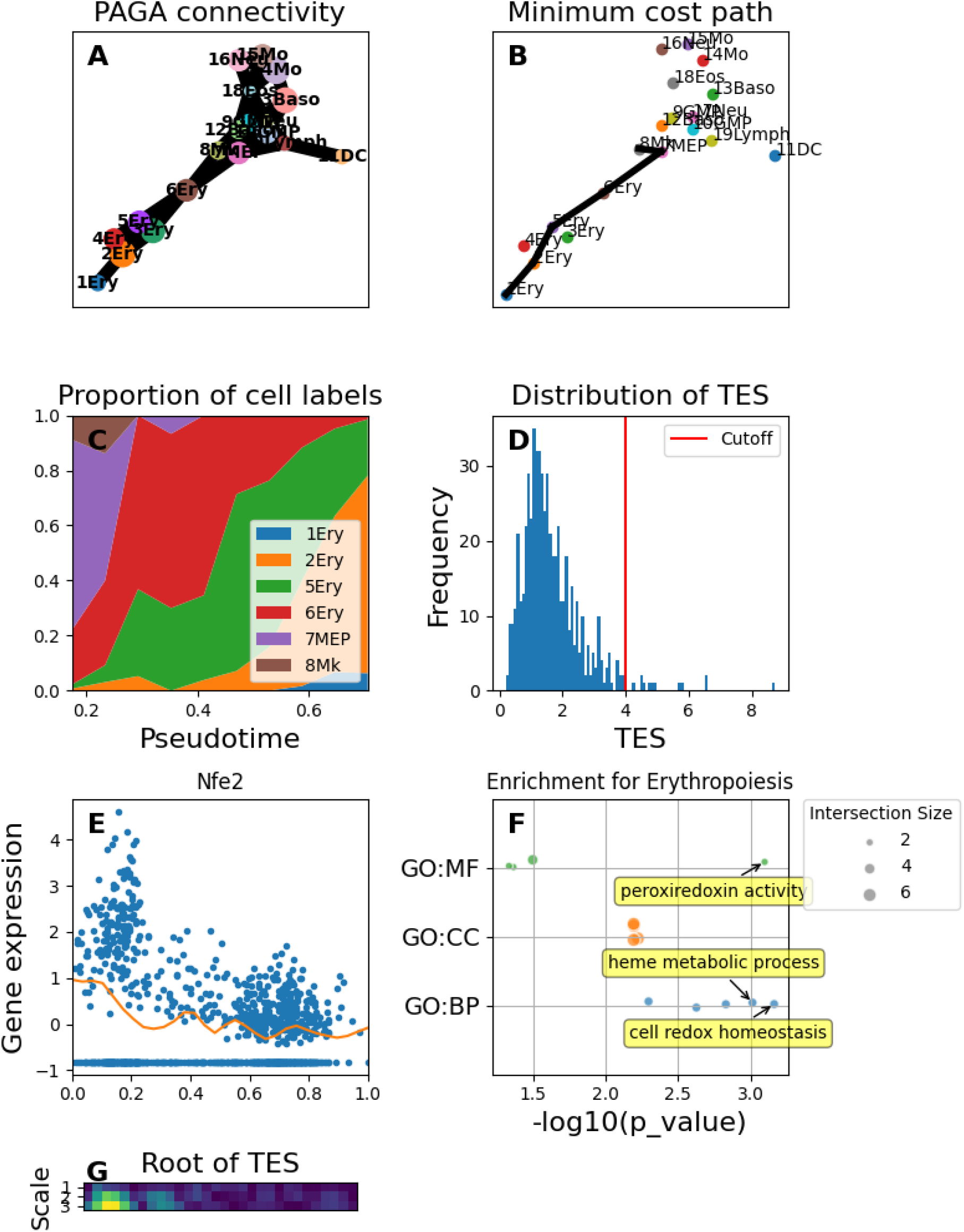
Pseudotime analysis of the hematopoiesis dataset, focusing on the erythropoiesis path. (A) Connectivity graph obtained by PAGA, thresholded to a minimum connectivity of 0.8. Labels are supplied with the dataset. (B) Minimum-cost path along the labels. Only cells belonging to these clusters are selected for processing. (C) Proportion of the cell labels across pseudotime. We see erythrocyte progenitors in the early parts of pseudotime get replaced by erythrocytes in later pseudotime. (D) After converting the gene expression to pseudotimecourses and scoring them to detect transient events, we obtain the distribution shown here. Genes scoring above the threshold are used for gene enrichment analysis. (E) Pseudotimecourse of Nfe2, a transcription factor necessary for erythropoiesis. (F) Functional gene enrichment analysis of the identified genes. A full list of the pathways can be found in Table 4. (G) The scale/position score for the pseudotimecourse, obtained by taking the absolute value of the product of the wavelet coefficients and their modified Z score. The square root is used to make the lower scores (blue-teal) more visible.

#### Monocyte to macrophage transition

Data published by Specht et al. for the SCoPE2 paper [12] consists of both transcriptomic and proteomic data for monocytes and macrophages. Monocytes differentiate into macrophages upon leaving the bloodstream and entering tissues [22]. The dataset offers some insight into the differences between omic layers in pseudotime. **Figure 4A** shows the distribution of cells in pseudotime when ordered using scRNA-seq data. The monocytes and macrophages are clearly differentiable, with few cells in between. The separation in pseudotime points to omic profiles that are distinct from one another. As such, it is not possible to extract a meaningful pseudotimecourse since most of the signal would be dictated by a handful of cells. In contrast, the proteomic data has the two cell types together in pseudotime (**Fig. 4B**), suggesting greater similarity in the proteomic profiles and a smoother transition between the two. Note that although DPT normalizes pseudotime values to be between 0 and 1, the distribution appears clipped because the 0.5-1.0 range consisted primarily of outliers (<3% of data). These were removed from subsequent analysis. Using the supplied metadata to identify cell type, we can see the transition of monocyte to macrophage along pseudotime (**Fig. 4C)**. A sample pseudotimecourse showing the expression of CAPG, a marker of macrophages, gradually increases along the obtained pseudotime (**Fig. 4D)**. Applying a TES threshold to obtain 130 genes, and clustering their pseudotimecourses using kmeans [23] (k=4) clustering, we obtain groups of genes with similar changes in expression (**Fig. 4E-I**). To examine whether the obtained clusters are biologically meaningful, we performed gene enrichment analysis and obtained the pathways shown in **Table 2-4** (cluster 0 had no significant enrichment). We note the previously-published relevance to monocyte/macrophage function or differentiation of AnxA2-p11 [24] complex, membrane raft organization [25], and S100 protein binding [26] in cluster 2 (**Table 3**), and L-malate dehydrogenase activity [27] for cluster 3 (**Table 4**).

**Table 2.** Functional gene enrichment pathways for the monocyte to macrophage dataset, cluster 1.

| source | native | name | p_value |
| --- | --- | --- | --- |
| GO:CC | GO:0070062 | extracellular exosome | 0.00663 |
| GO:CC | GO:1903561 | extracellular vesicle | 0.00731 |
| GO:CC | GO:0043230 | extracellular organelle | 0.00734 |
| GO:CC | GO:0065010 | extracellular membrane-bounded organelle | 0.00734 |
| GO:CC | GO:0005925 | focal adhesion | 0.01526 |
| GO:CC | GO:0030055 | cell-substrate junction | 0.01703 |
| GO:BP | GO:0051017 | actin filament bundle assembly | 0.03901 |
| GO:BP | GO:0061572 | actin filament bundle organization | 0.04198 |

**Table 3.** Functional gene enrichment pathways for the monocyte to macrophage dataset, cluster 2.

| source | native | name | p_value |
| --- | --- | --- | --- |
| GO:MF | GO:0048306 | calcium-dependent protein binding | 2.07E-06 |
| GO:CC | GO:0016363 | nuclear matrix | 1.62E-05 |
| GO:CC | GO:1990665 | **AnxA2-p11 complex** | 1.81E-05 |
| GO:CC | GO:0034399 | nuclear periphery | 3.23E-05 |
| GO:BP | GO:1903729 | regulation of plasma membrane organization | 4.91E-05 |
| GO:BP | GO:0010954 | positive regulation of protein processing | 0.00012 |
| GO:BP | GO:1903319 | positive regulation of protein maturation | 0.00016 |
| GO:BP | GO:0031579 | **membrane raft organization** | 0.00016 |
| GO:MF | GO:0005509 | calcium ion binding | 0.0004 |
| GO:BP | GO:1905686 | positive regulation of plasma membrane repair | 0.00109 |
| GO:MF | GO:0044548 | **S100 protein binding** | 0.00164 |
| GO:CC | GO:0062023 | collagen-containing extracellular matrix | 0.00193 |
| GO:BP | GO:0070613 | regulation of protein processing | 0.00237 |
| GO:BP | GO:1903317 | regulation of protein maturation | 0.00285 |
| GO:BP | GO:1905684 | regulation of plasma membrane repair | 0.0038 |
| GO:BP | GO:0010756 | positive regulation of plasminogen activation | 0.00506 |
| GO:CC | GO:0031012 | extracellular matrix | 0.00549 |
| GO:CC | GO:0030312 | external encapsulating structure | 0.00553 |
| GO:CC | GO:0070062 | extracellular exosome | 0.00599 |
| GO:CC | GO:1903561 | extracellular vesicle | 0.0064 |
| GO:CC | GO:0043230 | extracellular organelle | 0.00641 |
| GO:CC | GO:0065010 | extracellular membrane-bounded organelle | 0.00641 |
| GO:BP | GO:0001765 | membrane raft assembly | 0.00994 |
| GO:BP | GO:0010755 | regulation of plasminogen activation | 0.02761 |
| GO:CC | GO:0005576 | extracellular region | 0.02873 |
| GO:MF | GO:0008092 | cytoskeletal protein binding | 0.04821 |
| GO:BP | GO:0007009 | plasma membrane organization | 0.04933 |

**Table 4.** Functional gene enrichment pathways for the monocyte to macrophage dataset, cluster 2.

| source | native | name | p_value |
| --- | --- | --- | --- |
| GO:CC | GO:0005829 | cytosol | 5.34E-06 |
| GO:CC | GO:0031974 | membrane-enclosed lumen | 5.82E-05 |
| GO:CC | GO:0070013 | intracellular organelle lumen | 5.82E-05 |
| GO:CC | GO:0043233 | organelle lumen | 5.82E-05 |
| GO:CC | GO:0070062 | extracellular exosome | 6.63E-05 |
| GO:CC | GO:1903561 | extracellular vesicle | 7.96E-05 |
| GO:CC | GO:0065010 | extracellular membrane-bounded organelle | 8.02E-05 |
| GO:CC | GO:0043230 | extracellular organelle | 8.02E-05 |
| GO:CC | GO:0043227 | membrane-bounded organelle | 0.00012 |
| GO:CC | GO:0005737 | cytoplasm | 0.00014 |
| GO:MF | GO:0051082 | unfolded protein binding | 0.00064 |
| GO:CC | GO:0005654 | nucleoplasm | 0.00074 |
| GO:CC | GO:0005622 | intracellular anatomical structure | 0.0008 |
| GO:CC | GO:0001673 | male germ cell nucleus | 0.00363 |
| GO:CC | GO:0022626 | cytosolic ribosome | 0.00391 |
| GO:CC | GO:0031981 | nuclear lumen | 0.00432 |
| GO:CC | GO:0043226 | organelle | 0.00452 |
| GO:BP | GO:0006457 | protein folding | 0.00656 |
| GO:CC | GO:0043073 | germ cell nucleus | 0.00894 |
| GO:BP | GO:0002181 | cytoplasmic translation | 0.01392 |
| GO:CC | GO:0005615 | extracellular space | 0.01804 |
| GO:MF | GO:0044183 | protein folding chaperone | 0.02047 |
| GO:CC | GO:0043231 | intracellular membrane-bounded organelle | 0.02565 |
| GO:MF | GO:0030060 | **L-malate dehydrogenase (NAD+) activity** | 0.0298 |
| GO:CC | GO:0031982 | vesicle | 0.03543 |
| GO:CC | GO:0044391 | ribosomal subunit | 0.04131 |

**Figure 4.**
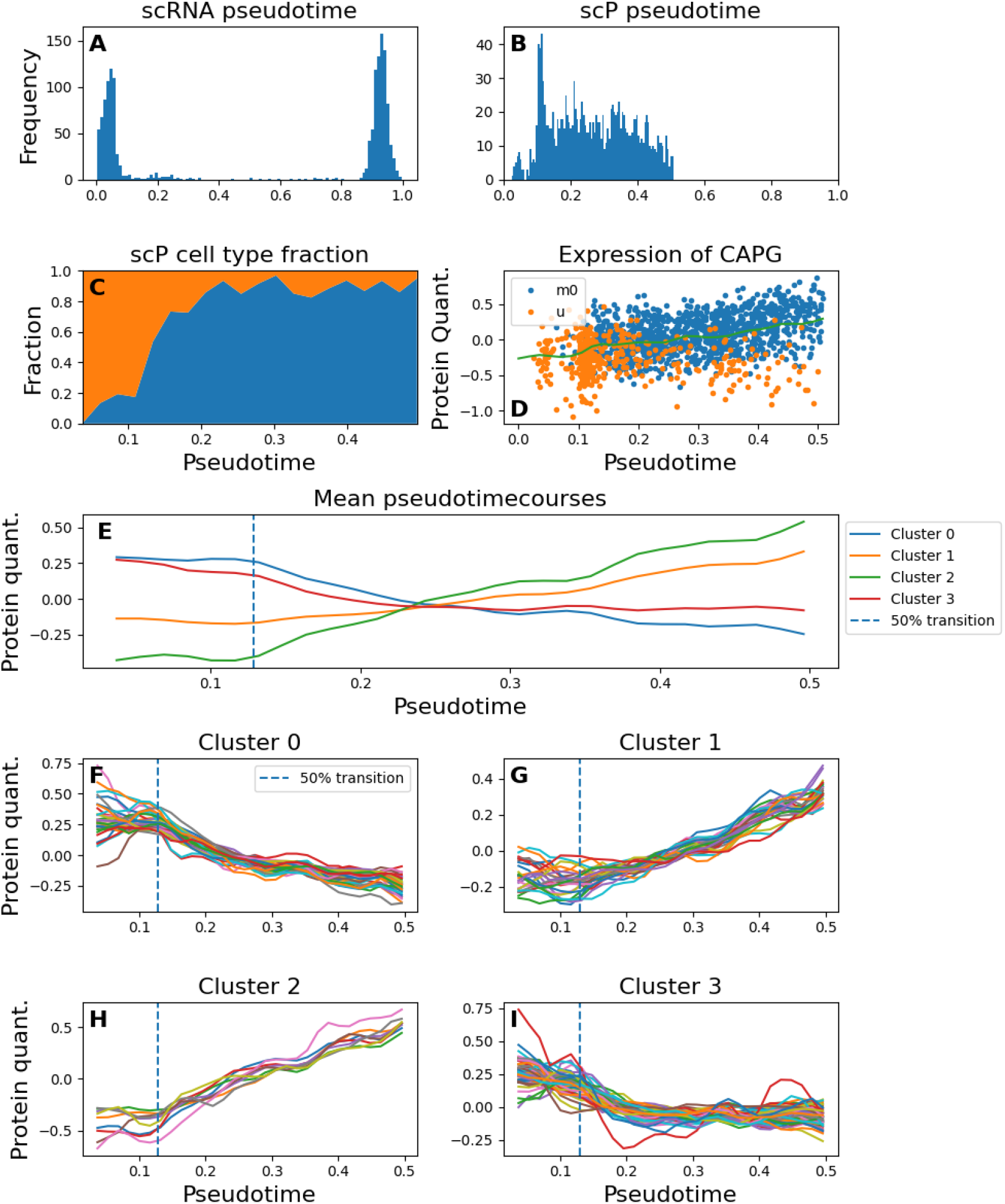
Pseudotime results for the scRNA and scP data for monocyte and macrophages published in the SCoPE2 paper. (A) Distribution of cell samples across pseudotime for the scRNA data. The wide gap in the distribution indicates large differences in the transcriptomic profiles of the cell types. (B) Distribution of cell samples across pseudotime for the scP data. The distribution shows no gaps, indicating that the protein profiles are similar between the cells. (C) Fraction of cell type in the scP data across pseudotime. The monocytes (orange) are grouped near the beginning of pseudotime, and the population fraction shifts towards macrophages (blue). (D) Sample pseudotimecourse for CAPG. (E) Mean pseudotimecourse for the four clusters of genes identified as having TEs. The clusters demonstrate different mean pseudotimecourses, but the means miss some of the TE seen in some of the individual traces. (F-I) Pseudotimecourses of each gene within a cluster.

We note that the proteins identified by scTransient were not limited to only those that were differentially expressed between the cell types (**Suppl. Fig. 1**, see Discussion).

#### Cell cycle

A similar analysis was performed using deep single cell proteomics data from human lung adenocarcinoma A549 cells produced by Bubis et al. [16]. One of the advantages stated in the study is the ability to study the dependency between a cell’s size and its cell cycle phase, making the data well-suited for evaluating our proposed method. Using default parameters for Leiden clustering, four clusters were detected for the dataset (**Fig. 5A**) that were not clearly associated with the proteome depth in those cells (**Fig 5B**). We identify the cell cycle phase using Scanpy’s *score_genes_cell_cycle* function that applies gene scoring as described in Seurat [28] by supplying known markers for S and G2/M phases, and labelling the rest as G1. The cell cycle showed coarse agreement with the Leiden clusters (**Fig 5C**). We set ordinal labels of 1-3 for the G1, S, and G2/M phases and used psupertime [6] to obtain pseudotime orderings. Proteins found to be significant in predicting cell cycle phase are shown in **Fig. 5D**, and plotting the cell phase as a function of pseudotime produces the expected transitions between phases (**Fig. 5E**). The trajectory of some well-known cell cycle proteins PCNA and UNG over pseudotime is consistent with their role in the S phase (**Suppl. Fig. 2**).

**Figure 5.**
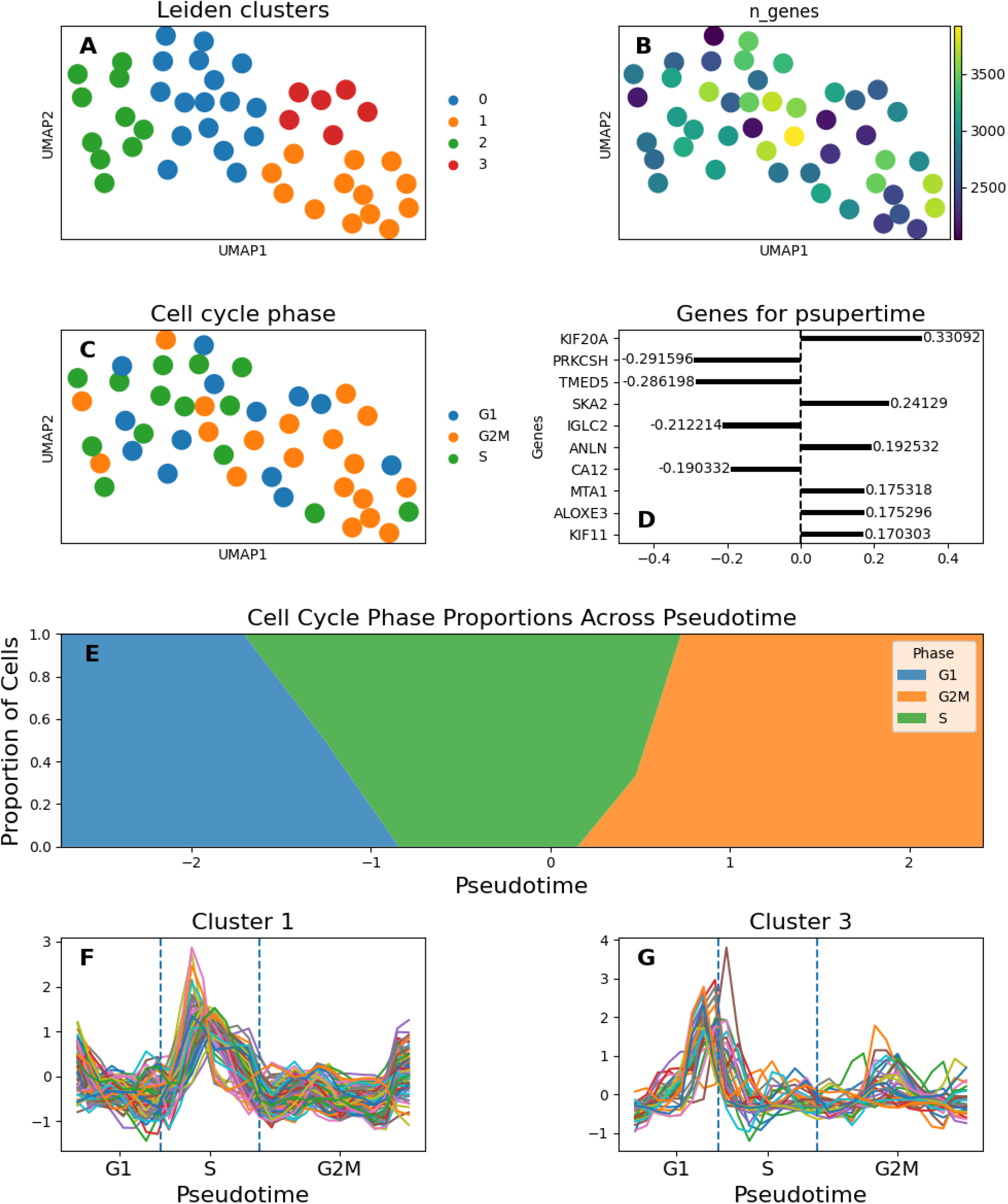
Analysis of the cell cycle dataset. (A) Cell phase proportions across pseudotime. After identifying the cell phase using phase biomarkers and ordering them using psupertime, we see a gradual transition across cell cycle phases. (B) Pseudotimecourse for cluster 1, with increased protein quantities for cells in the S-phase. The enriched pathways listed in Table 1 are consistent with DNA replication. (C) Pseudotimecourse for cluster 2, showing increased protein quantities for cells near the end of the G1 phase.

We apply windowing as previously described and use scTransient to identify TEs. Although no clear pattern is apparent in the highest-scoring genes, obtaining approximately 200 genes and clustering them using kmeans (k=4) clustering, we see two relevant patterns emerge. One cluster consists of genes (n=83) spiking in the area of pseudotime (**Fig. 5F**) related to the S-phase, with functional gene enrichment analysis using g:Profiler [20], [21] (**Table 5**) showing links to DNA replication, RNA processing, and nucleus-related activity. The second cluster consists of genes (n=32) genes spiking near the G1/S transition region (**Fig. 5G**), with gene enrichment showing genes related to extracellular space among others (**Table 6**). We note that there exists literature suggesting that components listed in **Table 6** are related to the cell cycle in different cell types, but we are unable to experimentally verify that this is the case for the dataset. Of the two remaining clusters, one showed no significant enriched pathways, and the other had pathways related to NADPH and extracellular components (Table 7). We note that for the cluster containing 83 proteins, 22 of these did not have previously published literature supporting a connection to the cell cycle (**Supplemental Table 1)**; some of the other 61 had links to altering the cell cycle in cancer cell lines. Experimentally validating the impact of these proteins is beyond the scope of the current paper.

**Table 5.** Significant pathways for the cluster 1 of the genes found to have transient events in the cell cycle dataset.

| source | native | name | p_value |
| --- | --- | --- | --- |
| GO:CC | GO:0005654 | nucleoplasm | 7.96E-12 |
| GO:CC | GO:0031981 | nuclear lumen | 3.52E-10 |
| GO:CC | GO:0005634 | nucleus | 2.35E-09 |
| GO:CC | GO:0031974 | membrane-enclosed lumen | 7.11E-09 |
| GO:CC | GO:0043233 | organelle lumen | 7.11E-09 |
| GO:CC | GO:0070013 | intracellular organelle lumen | 7.11E-09 |
| GO:MF | GO:0003723 | **RNA binding** | 6.66E-08 |
| GO:CC | GO:0043231 | intracellular membrane-bounded organelle | 1.19E-07 |
| GO:MF | GO:0140640 | **catalytic activity, acting on a nucleic acid** | 2.30E-07 |
| GO:CC | GO:0043232 | intracellular membraneless organelle | 3.95E-07 |
| GO:CC | GO:0043228 | membraneless organelle | 3.97E-07 |
| GO:CC | GO:0043229 | intracellular organelle | 7.75E-07 |
| GO:CC | GO:0005622 | intracellular anatomical structure | 8.78E-06 |
| GO:CC | GO:0043227 | membrane-bounded organelle | 1.40E-05 |
| GO:MF | GO:0003676 | **nucleic acid binding** | 1.42E-05 |
| GO:CC | GO:0043226 | organelle | 2.55E-05 |
| GO:BP | GO:0016072 | **rRNA metabolic process** | 0.000137 |
| GO:MF | GO:0008173 | **RNA methyltransferase activity** | 0.000142 |
| GO:MF | GO:0004386 | **helicase activity** | 0.000144 |
| GO:MF | GO:0140098 | **catalytic activity, acting on RNA** | 0.000341 |
| GO:CC | GO:0005730 | nucleolus | 0.000379 |
| GO:BP | GO:0006364 | **rRNA processing** | 0.000421 |
| GO:MF | GO:0097159 | organic cyclic compound binding | 0.000687 |
| GO:BP | GO:0042254 | **ribosome biogenesis** | 0.000749 |
| GO:BP | GO:0071840 | **cellular component organization or biogenesis** | 0.001483 |
| GO:BP | GO:0001510 | **RNA methylation** | 0.001618 |
| GO:CC | GO:0035770 | ribonucleoprotein granule | 0.001713 |
| GO:MF | GO:0005515 | protein binding | 0.005665 |
| GO:BP | GO:0090304 | **nucleic acid metabolic process** | 0.007752 |
| GO:BP | GO:0009451 | **RNA modification** | 0.009092 |
| GO:CC | GO:0036464 | cytoplasmic ribonucleoprotein granule | 0.010392 |
| GO:CC | GO:0030684 | preribosome | 0.011902 |
| GO:CC | GO:0140513 | nuclear protein-containing complex | 0.01326 |
| GO:CC | GO:0005829 | cytosol | 0.018899 |
| GO:BP | GO:0043414 | macromolecule methylation | 0.021186 |
| GO:BP | GO:0043170 | macromolecule metabolic process | 0.026077 |
| GO:BP | GO:0006260 | **DNA replication** | 0.026904 |
| GO:CC | GO:0098687 | **chromosomal region** | 0.027403 |
| GO:BP | GO:0006139 | nucleobase-containing compound metabolic process | 0.028182 |
| GO:CC | GO:0032991 | protein-containing complex | 0.031968 |
| GO:MF | GO:0008757 | S-adenosylmethionine-dependent methyltransferase activity | 0.033176 |
| GO:CC | GO:0005663 | **DNA replication factor C complex** | 0.033592 |
| GO:MF | GO:0140097 | **catalytic activity, acting on DNA** | 0.036054 |
| GO:CC | GO:0032040 | small-subunit processome | 0.037421 |
| GO:BP | GO:0007275 | multicellular organism development | 0.037504 |
| GO:CC | GO:0099080 | supramolecular complex | 0.041997 |
| GO:MF | GO:0016787 | hydrolase activity | 0.047276 |
| GO:BP | GO:0006401 | **RNA catabolic process** | 0.049419 |

**Table 6.** Significant pathways for the cluster 3 of the genes found to have transient events in the cell cycle dataset.

| source | native | name | p_value |
| --- | --- | --- | --- |
| GO:CC | GO:0070062 | extracellular exosome | 3.46E-05 |
| GO:CC | GO:1903561 | extracellular vesicle | 3.98E-05 |
| GO:CC | GO:0065010 | extracellular membrane-bounded organelle | 4.00E-05 |
| GO:CC | GO:0043230 | extracellular organelle | 4.00E-05 |
| GO:CC | GO:1990660 | calprotectin complex | 3.72E-04 |
| GO:BP | GO:0051238 | sequestering of metal ion | 5.03E-04 |
| GO:CC | GO:0031982 | vesicle | 5.39E-04 |
| GO:CC | GO:0005615 | extracellular space | 1.38E-03 |
| GO:BP | GO:0070488 | neutrophil aggregation | 3.28E-03 |
| GO:BP | GO:0032119 | sequestering of zinc ion | 3.28E-03 |
| GO:MF | GO:0035662 | Toll-like receptor 4 binding | 5.46E-03 |
| GO:BP | GO:0002523 | leukocyte migration involved in inflammatory response | 5.52E-03 |
| GO:CC | GO:0034774 | secretory granule lumen | 1.38E-02 |
| GO:CC | GO:0060205 | cytoplasmic vesicle lumen | 1.44E-02 |
| GO:CC | GO:0031983 | vesicle lumen | 1.47E-02 |
| GO:MF | GO:0050544 | arachidonate binding | 1.91E-02 |
| GO:MF | GO:0050543 | icosatetraenoic acid binding | 0.019062 |
| GO:MF | GO:0050542 | icosanoid binding | 0.025392 |
| GO:CC | GO:0005576 | extracellular region | 0.028535 |
| GO:MF | GO:0050786 | RAGE receptor binding | 0.04073 |
| GO:MF | GO:0030234 | enzyme regulator activity | 0.04203 |

**Table 7.** Significant pathways for the cluster 2 of the genes found to have transient events in the cell cycle dataset.

| source | native | name | p_value |
| --- | --- | --- | --- |
| GO:MF | GO:0004090 | carbonyl reductase (NADPH) activity | 1.79E-02 |
| GO:CC | GO:0070062 | extracellular exosome | 2.32E-02 |
| GO:CC | GO:1903561 | extracellular vesicle | 2.55E-02 |
| GO:CC | GO:0043230 | extracellular organelle | 2.56E-02 |
| GO:CC | GO:0065010 | extracellular membrane-bounded organelle | 2.56E-02 |
| GO:MF | GO:0004090 | carbonyl reductase (NADPH) activity | 1.79E-02 |
| GO:CC | GO:0070062 | extracellular exosome | 2.32E-02 |
| GO:CC | GO:1903561 | extracellular vesicle | 2.55E-02 |
| GO:CC | GO:0043230 | extracellular organelle | 2.56E-02 |
| GO:CC | GO:0065010 | extracellular membrane-bounded organelle | 2.56E-02 |

The cell cycle genes used to order the samples could plausibly be identified as having transient expression, which could limit the relevance of the method; we explore this in the discussion section.

## Discussion

### Synthetic Experiments

The first set of experiments involved the signal-to-noise ratio (SNR) and showed that for a given sample count, the TES of genes containing a TE improved with an increasing SNR (**Fig. 2C**). Increasing the sample count decreased the TES of noise genes, as expected. When the data is converted from samples along pseudotime to a signal, the averaging window considers points over a stretch of pseudotime; having additional points averages out the noise component and results in a flatter signal (which in turn reduces the Zmod component of the heuristic). Although the results shown in **Fig. 2C** might appear to suggest that a stronger signal can overcome low sample counts, the bimodal distribution shown in **Fig. 2E** confirms that this is not the case. For low-sample datasets, the improving score is driven by the iterations where samples happen to be located at the TE; experimentally, this would translate to hoping that one of your cells happened to be in the middle of the process being investigated. In that case, we would not expect detectability to improve beyond some SNR threshold, which is consistent with the results shown in **Fig. 2F**. The false negative rate is high for all signals when no signal is detectable (SNR=0), shows an initial decrease as some instances have samples land in the right region, then tapers off to a minimum that is dependent on the number of samples. Combined, these results confirm that using pseudotime ordering requires enough samples to derive reliable conclusions from the pseudotime signal. These results are based on multiple idealized conditions and should be treated as *extremely optimistic* if used to inform the design of biological experiments.

For biological events that could have multiple TEs, we showed that the proposed heuristic had a gradual decrease with an increasing number of events due to an overall decrease in the modified Z score (**Fig. 2G**). For the specified conditions, the overall score did not drop to the noise floor, but the combination of multiple conditions (lower SNR, fewer samples) might do so.

Lastly, transition time (sampling density during the TE) decreased the heuristic score, particularly for low-sample datasets (**Fig. 2H**). With fewer samples to describe the change in signal intensity, the peak in gene expression is less likely to be caught, which in turn decreases the wavelet coefficients and the overall score. The effect is most notable in low-sample datasets, where it drastically reduced the mean TES. There is a decreasing utility with the number of samples; even if a peak is not completely described, a transient change that can be reliably detected is not detected better with additional samples.

### Existing Datasets

#### Hematopoiesis

Using PAGA, we identified the connectivities between the clusters supplied with the hematopoietic dataset from Paul et al. [15] and found a differentiation trajectory from stem cells to erythrocytes. Converting the gene expression along the trajectory to a pseudotime signal, we were able to identify a set of genes that correspond to erythrocyte activity (namely peroxiredoxin activity, cell redox homeostasis, and heme metabolism, **Table 1**). The identified genes were identified primarily by high-scale (wide) wavelets at either the beginning or end of pseudotime, which describes a higher expression in one cluster of cells compared to clusters in other parts of pseudotime. Effectively, this amounts to examining the differential expression between the early parts of pseudotime and the rest. Some of the gene pseudotimecourses (e.g. Nfe2, **Fig. 3E**) showed multiple peaks throughout the pseudotimecourse. The results for this dataset are not sufficiently convincing to assert that TEs have been detected but show that information related to the biological process can nonetheless be detected.

#### Monocyte to macrophage transition

The results of the monocyte/macrophage dataset suggest that analyses downstream of pseudotime (such as this one) may be better suited for proteomic data over transcriptomic. The separation demonstrated in **Fig. 4A** for scRNA profiles indicate that the cell state transition is much faster with transcriptomics than proteomics.

Given the demonstrated effects of low sample counts and the reduced sampling density around TEs from the synthetic data experiments, we expect that scRNA-seq experiments would require a prohibitive number of samples to sufficiently characterize the transition for some processes. We note that the hematopoietic dataset also used scRNA-seq, so the benefit of proteomics over transcriptomics does not apply to *all* processes.

Using only the most significant genes did not immediately identify a small set of relevant genes but considering more genes and subsequently reducing their number by clustering them together produced plausible groups. The result suggests that there are potential refinements that could be made to the TES heuristic to better identify single-gene TEs. Without the additional experimental validation of multiple biological processes that are beyond the scope of this paper, it is not immediately possible to evaluate changes to scoring and whether they would be an improvement in performance.

As discussed for the hematopoiesis dataset, the proposed methods can select large-scale wavelets that would effectively correspond to examining differentially expressed (DE) genes, which would severely limit its relevance. We examined the top 100 DE genes between monocytes and macrophages as well as the genes with the top 100 TES. For clarity: DE genes are those that are expressed more in one group than the other; including decreased expression would give an identical set of genes for both groups. The genes identified as containing TEs did not entirely overlap with the DE genes of either the monocytes or macrophages (**Suppl. Fig. 1**). Half of the TE genes do not overlap with differentially expressed genes for either the monocyte or macrophages, demonstrating that the proposed method does not extract only redundant information. The trends visible in **Fig. 4E** don’t individually indicate the presence of TEs, but they all exhibit an inflection that corresponds to the 50% transition identified from cell labels. The cell type metadata was used to color **Figs. 4C-D** and determine the 50% transition but was not used to inform the pseudotime ordering. In this way, the pseudotimecourses of TE genes could be used to identify cell state transitions in datasets that don’t have labels for every sample.

#### Cell cycle

The cell cycle dataset produced by Bubis et al. [16] showed the most promising results. Cell cycle markers were used to order the cells, which might improve their downstream correspondence to known pathways compared to unsupervised pseudotime. Multiple TEs were detected that corresponded with the main processes of the S phase (**Fig. 5F, Table 5**) and the end of the G1 phase (**Fig. 5G, Table 6**). Detecting groups of genes demonstrating transient expression changes that align with the cell cycle phase shows the potential to use pseudotime to detect TEs in different biological processes. We note that the cell cycle dataset was acquired using more modern instrumentation compared to the monocyte/macrophage data (Astral, 2023 vs. Q-Exactive, 2013) that provided a better profile depth and a more reliable signal (higher SNR). It is unclear whether the detectability of TEs requires high-sensitivity instruments or if the difference in detectability is due to the biological processes being investigated.

Since the cell cycle phase was determined using a set of known cell cycle markers, it is reasonable to suspect that those same genes could be the ones identified by scTransient. Of the 97 genes known to be relevant to the cell cycle [29], 63 were present in the dataset. Clustering the pseudotimecourses of those genes into 4 groups using kmeans clustering [23], we show the groupings **Suppl. Fig. 3** and note that they do not have the increase in late G1 or throughout S as seen as in the clusters of the genes identified by scTransient (see also **Supplementary Table 2**).

## Conclusion

scTransient bridges a critical gap in single-cell analytics by turning trajectory inference from a descriptive ordering tool into a quantitative detector of fleeting regulatory programs. Across synthetic benchmarks and three biologically diverse public datasets, our wavelet-based Transient-Event Score consistently highlighted short-lived surges in gene or protein expression that map to process-specific pathways yet were overlooked by standard differential-expression approaches. The ability to recover erythropoiesis regulators, macrophage differentiation factors, and cell-cycle checkpoints underscores both the versatility of the method and the value of integrating proteomic as well as transcriptomic data. Because scTransient is deployed on the PSCS web platform, researchers can now explore transient events without local installation, iteratively tune parameters, and share reproducible workflows. Future extensions—including adaptive windowing, refined TES heuristics, and compatibility with multimodal atlases—promise to widen the landscape of discoverable biological “spikes.” We anticipate that charting these transient signals will reveal new regulatory nodes and therapeutic entry points in development, immunology, cancer, and beyond.

## Acknowledgements

This work was funded by the NIGMS (R35GM142502).

