## Supplementary figures for "Single-Cell Trajectory Inference for Detecting Transient Events in Biological Processes"

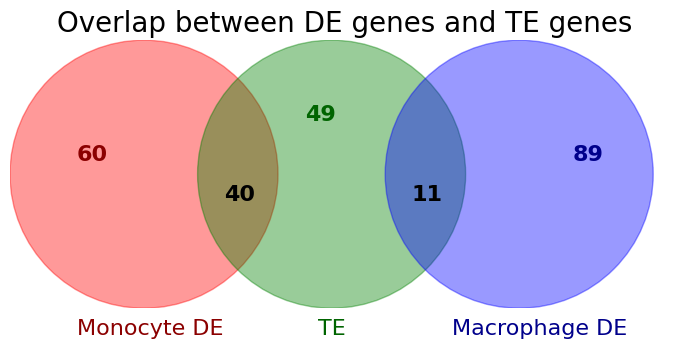

**Supplementary Figure 1.** Overlap between the top 100 differentially expressed genes and the top 100 genes identified to be related to a TE.

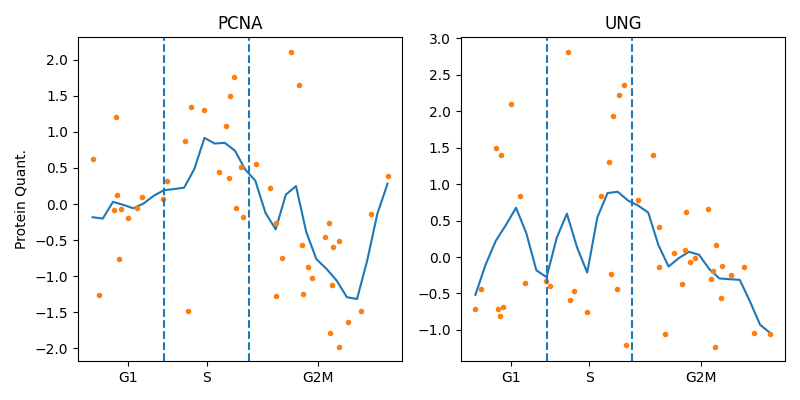

**Supplementary Figure 2.** Pseudotimecourses for two known cell cycle-regulating proteins. As expected, PCNA quantities peak in the S phase of the pseudotimecourse. The pseudotimecourse for UNG is noisier but shows a decline in G2/M similar to PCNA.

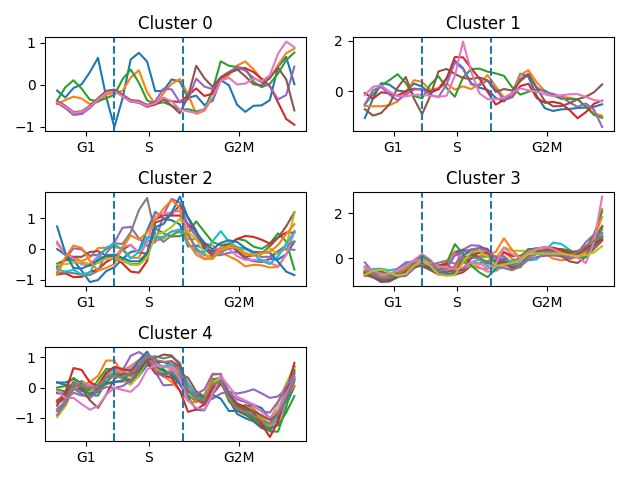

**Supplementary Figure 3.** Pseudotimecourse clustering of the proteins used to identify pseudotime ordering [[29]](https://www.zotero.org/google-docs/?fhtH92). The genes for each cluster can be found in supplemental table 2.

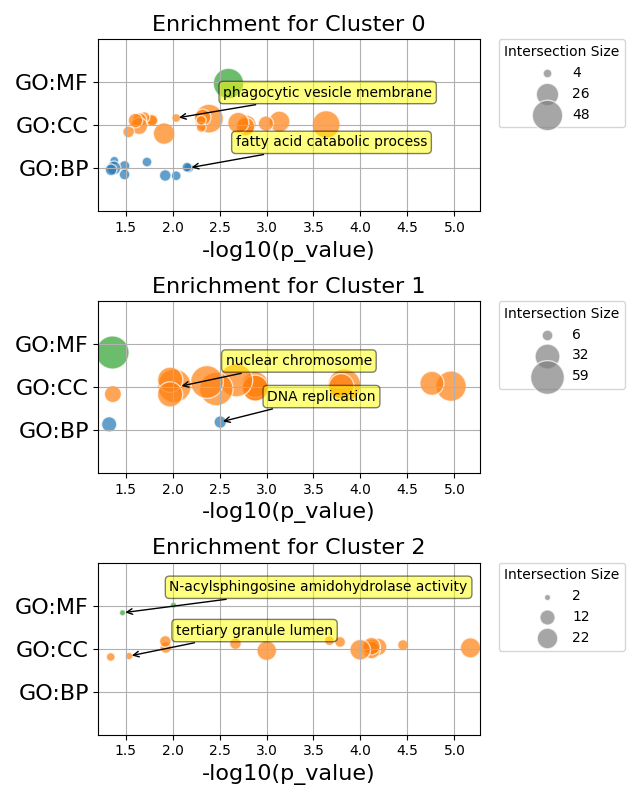

**Supplementary Figure 4.** Significant pathways found by gene enrichment [[20]](https://www.zotero.org/google-docs/?eRD3NF) for the clusters of genes identified for the cell cycle dataset [[16]](https://www.zotero.org/google-docs/?1fxlJP). Note that clusters 3 and 4 did not produce significant pathways.
